# ChIP-seq meta-analysis yields high quality training sets for enhancer classification

**DOI:** 10.1101/388934

**Authors:** Hana Imrichova, Stein Aerts

## Abstract

Genome-wide prediction of enhancers depends on high-quality positive and negative training sets. The use of ChIP-seq peaks as positive training data can be problematic due to high degrees of indirectly bound regions, and often poor overlap between experimental conditions.

Here we explore meta-analysis of ChIP-seq data to generate high-quality training data for enhancer modeling. Our method is based on rank aggregation and identifies a core set of directly bound regions per transcription factor, exploiting between five and twenty ChIP-seq data sets per factor. We applied this method to six different transcription factors, namely TP53, REST, SOX2, GRHL2, HIF1A and PPARG. Sequence analysis and modeling of recurrently bound enhancers yielded distinct enhancer features for the different factors, whereby binding sites of REST and TP53 are strongly determined by their motif; binding of GRHL2 and SOX2 is determined by nucleosome positioning; and binding of PPARG and HIF1A depends on other transcription factors. In conclusion, meta-analysis of ChIP-seq peaks, and centering on motifs, allowed discovering new properties of transcription factor binding.

## Introduction

Enhancers are “docking stations” for transcription factors (TFs). TFs bind to enhancers at specific DNA recognition sequences and regulate the expression of their target genes. In some cases, one TF is sufficient to bind to the DNA via its recognition motif and regulate gene expression. This is the case, for example, for TP53 (Sammons et al., 2015; Verfaillie et al., 2016) that binds to DNA as a tetramer and activates its target genes without additional TFs. However, in most cases, specific combinations of TFs bind coordinately to enhancers to achieve spatiotemporal control of gene expression, and consequently of the behavior, morphology and function of a cell (Lee and Young, 2013; Shlyueva et al., 2014). Such combinatorial binding of multiple TFs can either occur cooperatively or hierarchically (one TF binds as pioneer) (Reiter et al., 2017). Although intensive research has been done in the field of regulatory genomics in past years, several challenging questions remain unresolved. A key problem is the low accuracy of computational methods to predict functional TF binding sites and enhancers (Wasserman and Sandelin, 2004). Indeed, current methods have difficulties to discriminate bona fide from false positive sites, or TF-bound from TF-unbound sites (Aerts et al., 2005; Frith et al., 2003; Heinz et al., 2010; Medina-Rivera et al., 2015). To overcome the challenge of enhancer and TF binding site prediction, machine-learning methods are often used. When applied to enhancers, positive training sets of similar enhancers (i.e., enhancers with similar expression pattern) are usually contrasted against other enhancers or negative sequences, and have yielded satisfactory cross-validation results in multiple systems (Alipanahi et al., 2015; Lee et al., 2011; Svetlichnyy et al., 2015). However, classifiers that are trained on TF ChIP-seq peaks usually have very low performance. This was recently illustrated by the ENCODE-DREAM challenge (https://www.synapse.org/#!Synapse:syn6131484/wiki/402026). This is thought to be due to, at least partly, the low quality of ChIP-seq peaks, which can contain both direct and indirect binding, as well as phantom peaks resulting from cross-linking artifacts (Baranello et al., 2016; Keren and Segal, 2013; Wang et al., 2012).

To address this problem, and to investigate whether ChIP-seq peaks can be used to train reliable classifiers, we perform an in-depth analysis of ChIP-seq peaks, and introduce a meta-analysis framework to identify high-quality direct binding sites. Particularly, we build better training data sets for enhancer modeling by applying two different types of meta-analysis to panels of publicly available ChIP-seq data sets, obtained for the same TF but in different conditions or cell types (**Supplemental Table S1**). We investigate the use of threshold-free approaches, to allow for recurrent low-affinity binding; and the use of motif-guided signal detection. Overall, we find a surprisingly high concordance of *direct* TF binding across different cell types, tissues and conditions. We exploit this to define core sets of preserved functional binding sites to discover new enhancer features. Using machine learning models, we then evaluate these enhancer features and identify distinct TF binding modalities.

## Methods

### TF ChIP-seq datasets

TF ChIP-seq peaks were downloaded from ChIP-Atlas (http://chip-atlas.org) and quality-controlled by motif discovery using i-cisTarget (Imrichová et al., 2015), to ascertain that the motif of the ChIP’ped TF is enriched among the peaks. Subsequently, high-quality TF ChIP-seq datasets were downloaded as raw sequence reads (**Supplementary Table S1**). Reads were mapped to the reference genome (hg19-Gencode v18) using Bowtie2 2.2.6, with --*local* and --*sensitive-local* parameters. Reads with mapping quality lower 4 were filtered out.

### Selection of the candidate binding sites using genome-wide TF motif prediction

chromHMM method (Ernst and Kellis, 2012) was applied on ChIP-seq data per each TF separately to identify chromatin states. The parameter *-binsize 200* was used (default) and number of state 20 was specified. i-cisTarget (Imrichová et al., 2015), with a large motif collection including 18832 motifs, was applied on each chromHMM state to identify enriched motifs. The motif annotated for the ChIP’ped TF with a high enrichment score in the recurrent chromHMM state was selected (Figure 1, Figure 2c). Subsequently, candidate regulatory regions (as defined before (Imrichová et al., 2015)) were scored by Cluster-Buster (Frith et al., 2003) to obtain motif-cluster score for the selected motif. Moreover, the scoring was performed across genomes of 10 species, as described previously (Imrichová et al., 2015). This allowed us to predict genome-wide the top 20000 candidate regulatory regions with the highest motif-cluster scores for the selected motif across 10 genomes (Figure 2c,d).

**Figure 1.**
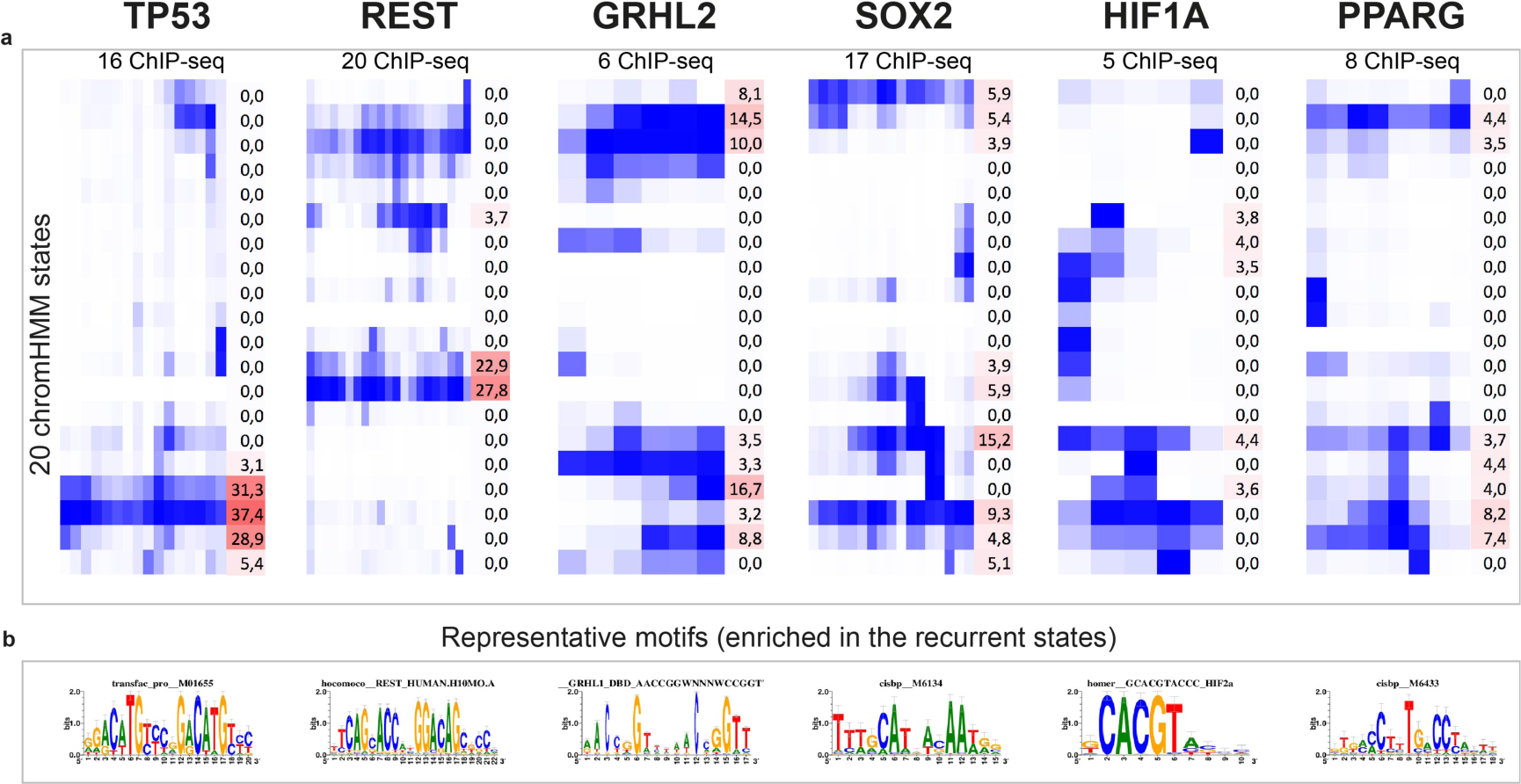
Meta-analysis by region clustering. **a)** Six TFs were selected for a meta-analysis by chromHMM (blue-white heatmap representing 20 chromHMM states), followed by motif discovery in each cluster/state (red-white heatmap representing the highest enrichment score of the motif of the ChIP’ped TF in the chromHMM state). **b)** Logos of the representative TF motifs. These motifs are annotated for the ChIP’ped TFs and were found highly enriched in the recurrent chromHMM states (i.e. these motifs are bound across different TF ChIP-seq experiments).

**Figure 2.**
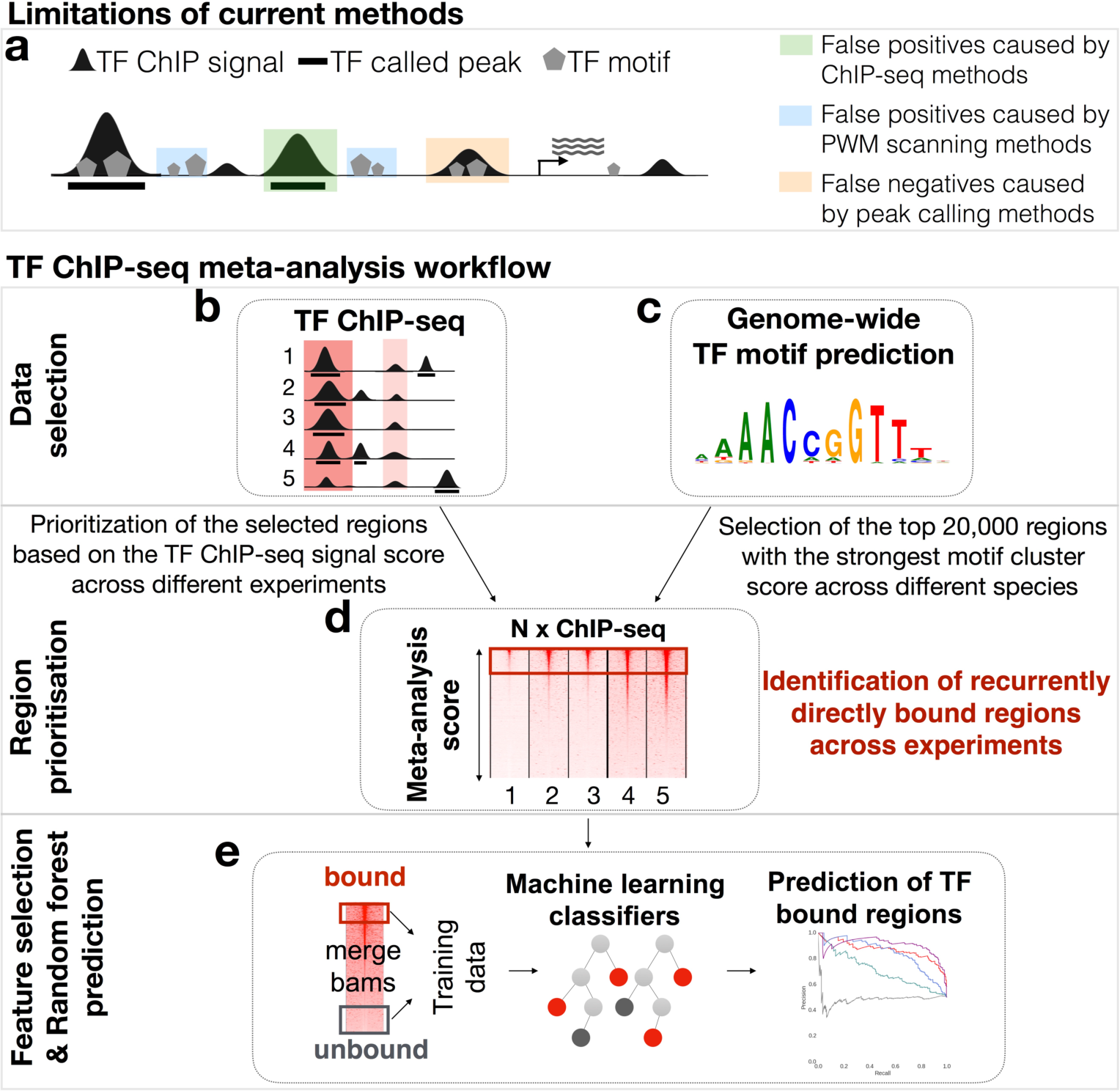
Workflow of the TF ChIP-seq meta-analysis. **(a)** Cartoon representing the limitations of existing computational and experimental methods. **(b-c)** ChIP-seq data sets and the representative motif are combined (the representative motif was obtained from the recurrent chromHMM states, see methods). **(d)** Region prioritization: the top 20.000 regulatory regions are selected based on motif cluster score of the representative motif across 10 mammalian genomes. The regions are re-scored in human genome (hg19) by Cluster-Buster and centered on the strongest motif. Subsequently, OrderStatistics is applied on TF ChIP-seq experiments to aggregate the signal information on the pre-selected candidate regulatory regions, hence prioritize the regions. The regions ranked at the top of the ChIP meta-analysis ranking are considered as a core set of bound regions persistent across different samples and conditions. **(e)** The bound and unbound regions are used to investigate TF binding modalities and to identify enhancer features to train machine learning classifiers.

Next, the selected 20000 regulatory regions that have strong motif cluster score for the specific motif across 10 species were re-scored by Cluster-Buster (Frith et al., 2003) (function *cbust*, using parameter *-m 0*) in human genome (hg19) to keep only regions with any score for the single motif in human. Finally, the regions were centered on the motif with the highest score detected by Cluster-Buster and extended to 500 bp upstream and 500 bp downstream from this motif. This procedure yielded a set of pre-selected regulatory regions per each TF.

### TF ChIP-seq meta-analysis using *OrderStatistics*

TF ChIP-seq regulatory data generated on different cell types under different conditions were used to prioritize the pre-selected regulatory regions using aggregation score which represents robustness of TF binding across different experiments, without peak calling threshold limitation. The coverage of TF ChIP-seq data on the pre-selected candidate regions with the match TF motif in hg19 was computed using function *coverageBed* from BEDTools v2.26.0. The regions were ranked based on the coverage score and subsequently the rankings across different samples and conditions were combined using *OrderStatistics* (Aerts et al., 2006; Stuart et al., 2003), into final meta-ranking per TF (Figure 2d, Figure 3b).

**Figure 3.**
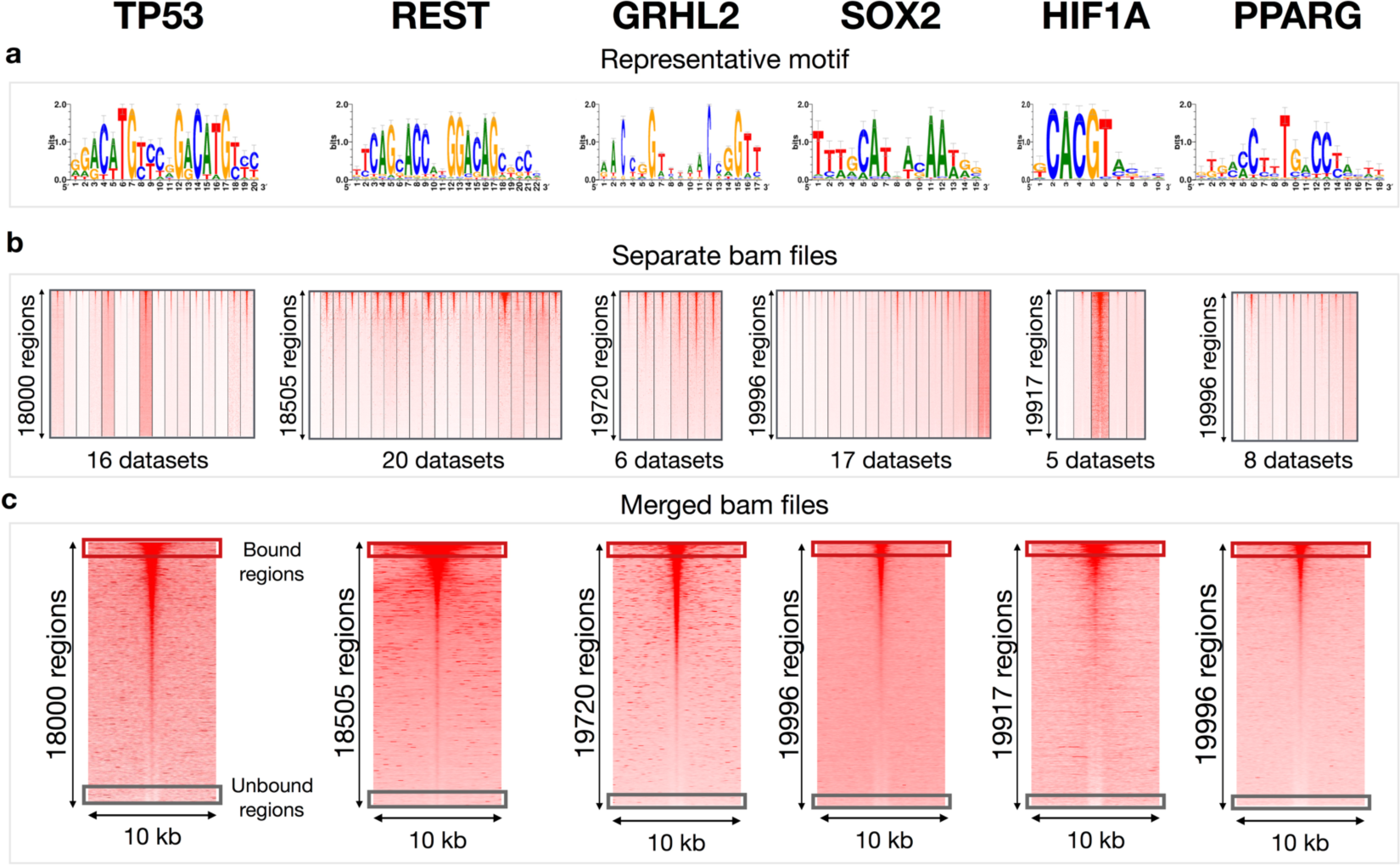
ChIP-seq meta-analysis by rank aggregation. **(a)** Logos of representative TF-binding motifs. **(b)** SeqMiner heatmaps representing TF ChIP-seq datasets on the selected regions ordered according to the TF ChIP-seq meta-analysis score. The number of pre-selected *cis*-regulatory regions where the representative motif was detected in the human sequence (hg19) is shown on the left. **(c)** SeqMiner heatmaps representing the signal of merged bam files from b) at the regions ordered according to the TF ChIP-seq meta-analysis score.

### seqMINER

seqMiner heatmaps were generated on the regulatory regions centered to the representative TF motif with the highest Cluster-Buster motif score. The seqMiner window 5000 bp upstream and downstream was used for visualization. The regions were ordered according to the TF ChIP meta-analysis score. As an input, bam files of reads with mapping quality >=4 were used (Figure 3b). To visualize a combined signal on the meta-ranked regions, bam files of all experiments per each TF were merged (generated by *merge* function from Samtools version 1.3 (Li et al., 2009)) (Figure 3c).

### Motif scores, GC content and prediction of nucleosome occupancy plots

The bound and unbound sequences (represented by the top and bottom 1000 regions based on TF ChIP-seq meta-analysis) were compared for motif score, GC content and nucleosome occupancy.

The highest Cluster-buster motif score per region was used to visualize the difference in motif strength between the bound and unbound regions. The statistical significance was computed by Wilcoxon rank sum test (two-sided) in R version 3.2.2.

The GC content information was calculated in bound and unbound sequences which represents occurrence of G and C nucleotides divided by the total number of base pairs. The GC fraction is visualized for the space 2 kb around the strongest motif.

For nucleosome occupancy, a file including genome-wide nucleosomes predictions (average occupancy, hg18) was downloaded from https://genie.weizmann.ac.il/software/nucleo_genomes.html (Kaplan et al., 2009). Liftover tool (Meyer et al., 2013) was used to map bound and unbound regions from hg19 to hg18 genome. Finally, the nucleosome preferred occupancy score was visualized in space 4 kb around the strongest motif.

### Machine learning prediction

Random forest models (Svetlichnyy et al., 2015) were trained and evaluated (the top and bottom 1000 regions based on TF ChIP-seq meta-analysis per each TF are used as positives and negatives, respectively, where the regions are centered to the strongest motif and extended to 500 bp upstream and 500 bp downstream from the motif). Each classifier consists of 151 decision trees. The performance of random forest models was evaluated by Precision Recall and generating the AUC values, which was calculated on left out data. Specifically, 80% of positive and negative sequences was used for training the models, while 20% of positive and negative sequences was used for testing.

Different combinations of features were tested: M1 model represents a model, where only one PWM annotated for the specific TF is used for training, M6 represents a model with six PWMs annotated for the specific TF (**Supplementary Figure S1**), M1_GC and M6_GC represent models where GC content information is used together with one and six motifs annotated for the specific TF, respectively.

Motif scores per region were obtained by Cluster-Buster, calculated on the full length of the sequence (1 kb around the motif). The GC fraction was calculated in the space 150 bp upstream and 150 bp downstream of the motif, which represents occurrence of G and C nucleotides divided by the total number of base pairs, and average value was used as the feature for Random forest models.

### DNase I hypersensitivity sites (DHS)

124 DHS experiments performed on different cell types and conditions were downloaded from ENCODE consortium database (ENCODE Project Consortium, 2012). The following data format was used: ENCODE broadPeak files and bam files. In case that more replicates were available, only the replicate 1 was used.

### Radar plot to represent the feature importance

The importance of four features (nucleosome occupancy, GC content, motif score by *cbust*, length of DHS peaks) was visualized using *radarchart* from R/Bioconductor package ‘*fmsb’*. For the first 3 features, the six studied TFs were assigned by a score on a scale from 1 (lowest importance) to 6 (highest importance) according to delta score between bound/unbound regions for the specific feature. The score for DHS length was assigned based on the ranking of TFs according to the median length of overlapping DHS peaks with the bound regions, where score 1 represents the broadest peaks and 6 represents the narrowest peaks.

## Results

### ChIP-seq meta-analysis using clustering techniques

We generated a focused compendium of 72 TF ChIP-seq data sets representing six different TFs, namely TP53, REST, GRHL2, HIF1A, SOX2, and PPARG. For each of these TFs, our compendium contains between 5 and 20 different experiments. All data sets were quality-controlled by motif discovery, to ascertain that the motif of the ChIP’ped TF is enriched among the peaks (**Supplementary Tables S2-S7**). Such a quality control step is often applied (Yan et al., 2013). In a first attempt to explore different modes of TF binding specificity, and avoiding ChIP-seq peak thresholds, we applied a clustering method based on Hidden Markov Models (chromHMM (Ernst and Kellis, 2012)). For each of the six TFs, chromHMM detects multiple states, but for each factor at least one state was found with recurrent TF binding signal (Figure 1a). Next, we analyzed each cluster of regions (i.e., each state) with i-cisTarget, a motif discovery method that uses a large collection of position weight matrices (Imrichová et al., 2015) to assess which of the states is associated with an enrichment of a motif that is annotated for the ChIP’ped TF.

For TP53 and REST, we observed the strongest and most significant enrichment of their motif in the recurrently bound states (Normalized enrichment score (NES) = 37.4 for TP53; and 27.8 for REST). Interestingly, the recurrent ChIP-seq signal of these TFs, that is associated with the match motif, is represented by only a few chromHMM states. For the remaining four TFs, the ChIP-seq signal is clustered into more heterogeneous chromatin states. The enrichment of the TF motif is again strongest in the recurrently bound regions, but for these factors, their motif is also enriched (although to a lesser extent) in several sample-specific states. For GRHL2 and SOX2, we obtained the NES = 14.5 and NES = 9.3, respectively, for the recurrent state. Of the six tested TFs, HIF1A and PPARG show the lowest motif enrichment scores (NES = 4.4 and NES = 7.4, respectively) (Figure 1). This analysis shows a wide diversity of binding heterogeneity, illustrating that using ChIP-seq peaks for training a classifier is not straightforward. To resolve this, we will introduce in the next section an alternative strategy for ChIP-seq meta-analysis, focused on directly bound regions (i.e., having a TF motif instance).

### Motif-centered ChIP-seq meta-analysis using rank aggregation

Our chromHMM clustering analysis of ChIP-seq data sets yielded a representative motif for each TF (Figure 1b), in the state representing recurrent binding of the TF across conditions. We used this representative motif in subsequent analyses to focus on directly bound regions. The workflow of our meta-analysis is depicted in Figure 2. For each of the six TFs, we created a set of 20.000 candidate binding sites, predicted across the genome (Figure 2 and Methods). We reasoned that, since most ChIP-seq data sets yield a few thousand peaks, the top 20.000 binding sites would contain bound as well as unbound sites. Next, we ranked these 20.000 regions based on the ChIP-seq signal (the raw read depth, thus again without ChIP-seq peak calling thresholds), for each TF ChIP-seq data set separately.

We applied this technique to the 72 TF ChIP-seq datasets, for the six selected TFs. This is followed by a rank-aggregation meta-analysis, *OrderStatistics* (Aerts et al., 2006; Stuart et al., 2003), where the final ranking contains regions that are most strongly bound across conditions. This resulted in a remarkable preservation of binding (Figure 3b), for each of the six TFs. From the top-scoring regions in this meta-ranking, we defined a core set of bound regions persistent across different samples and conditions, while those at the bottom are considered as unbound regions (Figure 3b,c). These bound and unbound regions will be used below to discover differences in binding preferences between studied TFs and to learn relevant sequence features that were further validated by machine learning predictions.

### Analysis of bound versus unbound sites

When comparing the bound and unbound regions (represented here by the top and bottom 1000 regions based on the TF ChIP-seq meta-ranking), we observed interesting differences between the TFs. Firstly, for TP53 and REST we found a highly significant difference in motif scores between bound and unbound (pval = 7.83*10^-191^ and pval = 6.27*10^-282^, respectively); while HIF1A and PPARG show similar motif scores whether the sequence is bound or not (pval = 2.56*10^-15^ and pval = 1.76*10^-22^, respectively) (Figure 4a). Thus, for TP53 and REST the motif score is highly predictive for their actual binding, while for HIF1A and PPARG the motif score is not immediately predictive. GRHL2 and SOX2 are intermediate, as they show a high significant difference in motif score between bound and unbound sites (pval = 1.79*10^-94^ and pval = 2.05*10^-67^, respectively), but not as strong as TP53 and REST.

**Figure 4.**
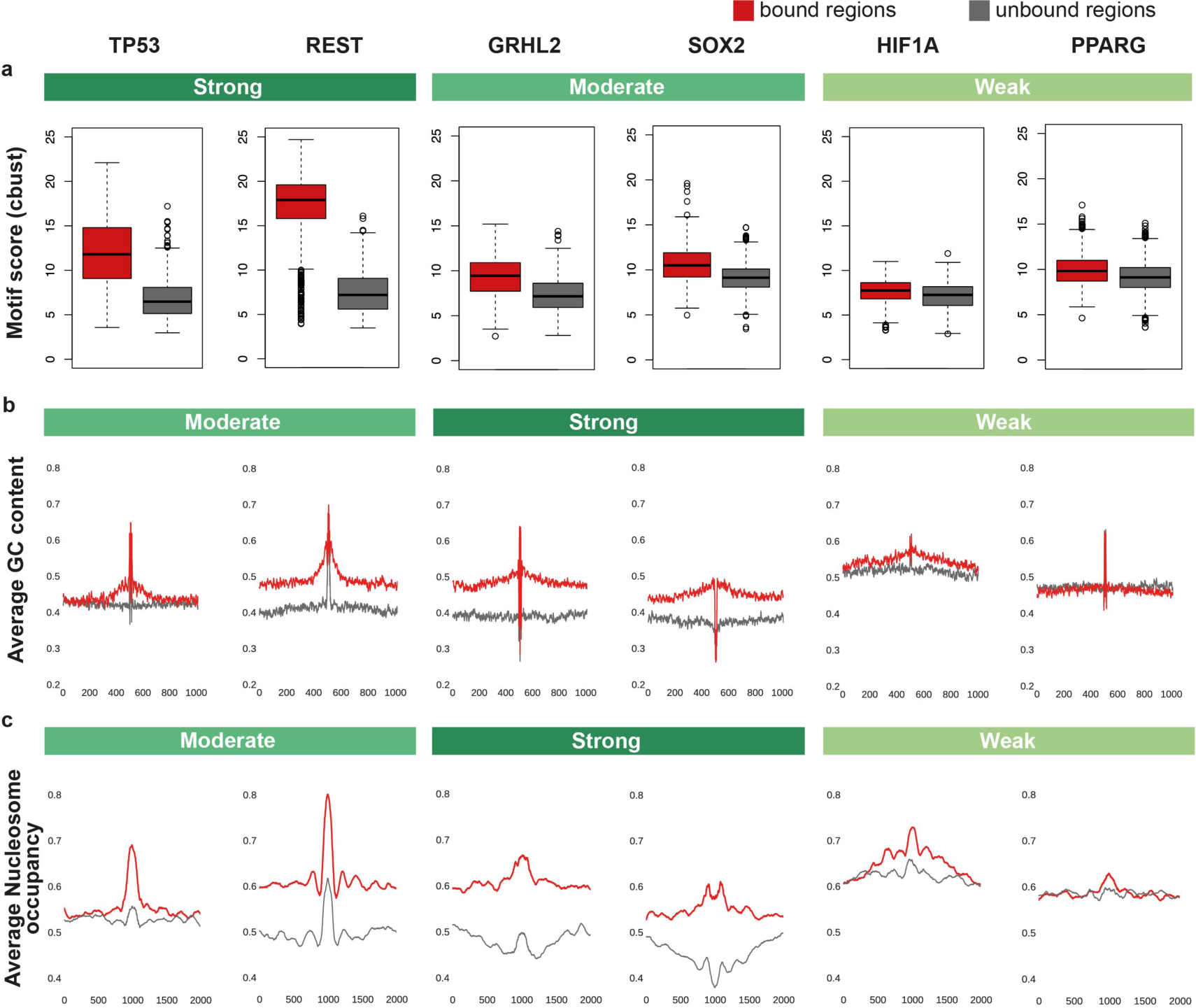
Differences between TFs when comparing bound and unbound regions. **a)** Motif score calculated by Cluster-Buster (cbust). **b)** Average GC content visualized for the sequence 2 kb around the strongest motif. **c)** Average nucleosome occupancy score visualized for the sequence 4 kb around the strongest motif.

Next, we compared other sequence features in the flanking sequences around the motif, between bound and unbound regions. A strong difference in GC content was detected for GRHL2 and SOX2, where the bound regions have higher GC content compared to their unbound counterparts (Figure 4b). Vice-versa, almost no difference in GC content was observed for HIF1A and PPARG.

Next, we examined whether differences in nucleosome preference could be detected between bound and unbound sites. SOX2 and GRHL2 emerge as factors that bind to regions with a strong nucleosome affinity (Figure 4c). This is a feature that is related to pioneering TFs that are able to detect their binding sites in silent/repressed chromatin. Their binding causes nucleosome displacement, which allows other TFs to bind and initiate enhancer activation (Jacobs et al., 2018; Soufi et al., 2015; Zaret and Mango, 2016).

### Random Forest discriminates between bound and unbound sites

Having discovered several motifs and sequence features for bound versus unbound regions, we exploited this information to train Random Forest models to discriminate bound from unbound TF motif instances. We tested several models, each time including different features for each TF to identify the most important features per TF. Interestingly, this revealed that for REST and TP53 a single motif (M1) is already sufficient to predict binding with high accuracy (Precision-Recall AUC (AUPR) = 0.97 and 0.87, respectively), suggesting very high motif dependence of these factors (Figure 5). For TP53, this result agrees with previously published results (Verfaillie et al., 2016). Predictions for the other TFs range from 0.49 (for HIF1A) to 0.71 (for GRHL2) and can be significantly improved by including multiple motifs of the same TF (M6) (**Supplementary Figure S1**), yielding performance increases by 13-15%. For SOX2, GRHL2, HIF1A and PPARG we observed that the GC content is very informative to predict binding, increasing the AUPR by 7-16% (M1_GC). Interestingly, using a combination of motifs and GC content (M6_GC) yielded even higher performances between 0.686 and 0.995 (for HIF1A and REST, respectively). Nucleosome affinity did not increase this performance further, which could be due to the correlated/confounding nature of GC content and nucleosome preference. Overall, the best performances were obtained for REST, TP53 and GRHL2, while additional information like co-factor motifs are needed for the other factors (for both HIF1A and PPARG the AUPR increases by 10% when co-factor motifs are included in the models, although the overall performance remains low (**Supplementary Figure S2**)).

**Figure 5.**
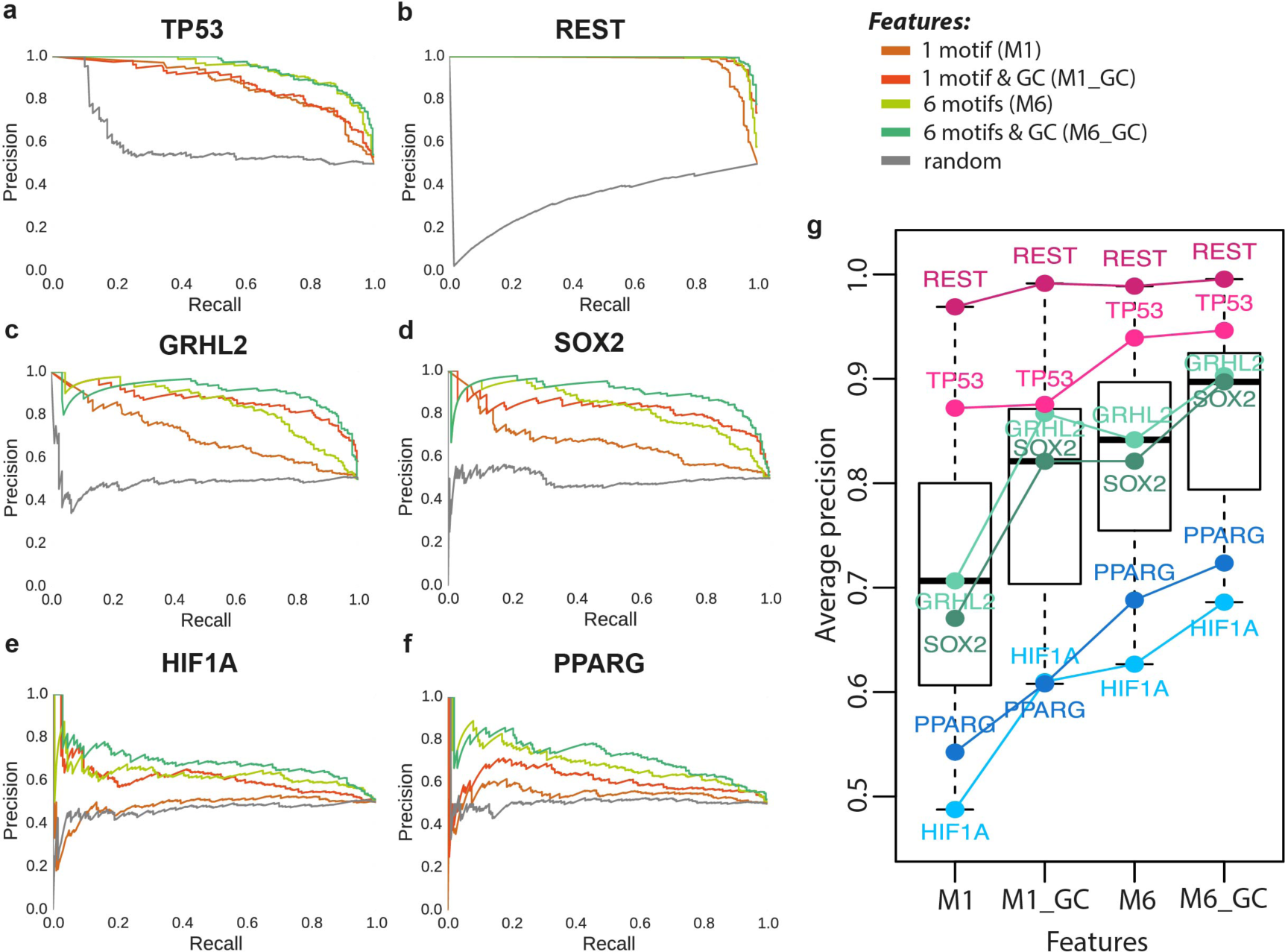
Random forest predictions of TF binding sites. **(a-f)** The plots represent Precision-Recall curves assessing the performance of a Random Forest classifier to discriminate between bound and unbound regions. The most accurate distinction across all TFs is achieved by a model using a combination of six motifs annotated for the specific TF together with GC fraction (green curve). **g)** Representation of the average precision-recall for each combination of features used for training the model.

### Binding of solitary factors results in narrow accessibility peaks

We identified two TFs, namely TP53 and REST, with very strong motif dependence, suggesting that they are solitary factors with unsophisticated enhancer models (Verfaillie et al., 2016). We reasoned that such solitary binding may be reflected by the open chromatin pattern of the bound regions. To test this, we analyzed publicly available DHS data (DNaseI-seq data from ENCODE (ENCODE Project Consortium, 2012)). First, we focused on the REST TF, a known repressor of neuronal genes in non-neuronal cells (Chong et al., 1995). We selected neuronal and non-neuronal cell lines for which DHS data is available and compared the signal at the REST core regions (predicted by ChIP-seq meta-analysis) between these two groups. Interestingly, we observed a sharp DHS signal on the REST core set regions in the non-neuronal samples (the samples where the REST is expected to repress the neuronal genes), suggesting that these sites are bound by a TF (**Supplementary Figure S3**, blue). Vice-versa, the same regions are depleted by the DNaseI signal in neuronal samples, suggesting that the regions are not bound by REST repressor or another TF (the neuronal genes are not supposed to be repressed in these samples).

Importantly, the sharp DHS signal suggests that REST acts alone, as a “solitary” TF and does not need another TFs to co-bind. We next investigated whether such a DHS pattern is also observed for the other TFs. As expected, TP53 shows the second most narrow DHS peaks (Figure 6), while the other non-solitary TFs are associated with a moderate (SOX2 and GRHL2) to high width (HIF1A and PPARG) of open chromatin peaks. This finding suggests a general trend that strong motif dependence is anti-correlated with the width of open chromatin regions (Figure 6).

**Figure 6.**
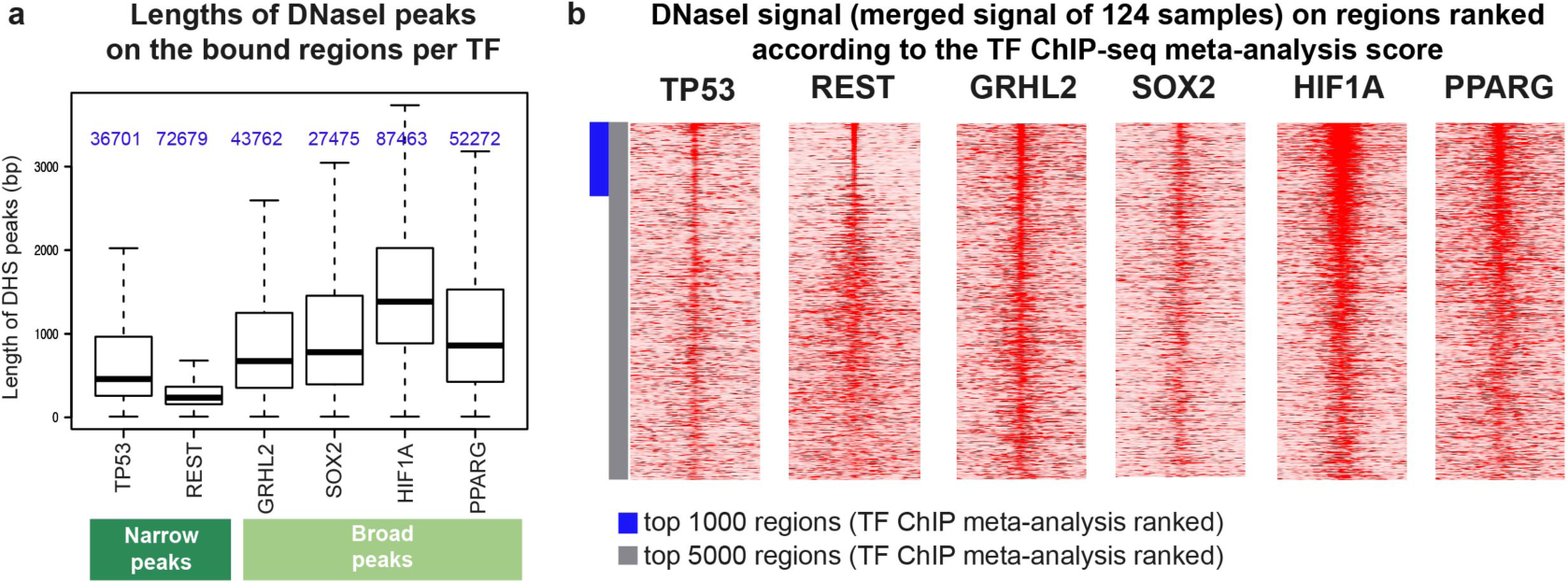
DNaseI signal mapped on the core set of bound regions. **a)** The length of DNaseI peaks (from 124 samples) that overlap the bound regions. **b)** The merged DNaseI signal of 124 samples on regions ranked according to the TF ChIP-seq meta-analysis score.

## Discussion

To build better training data sets for enhancer modeling, we apply two different types of meta-analysis to panels of publicly available ChIP-seq data sets, obtained for the same TF but in different conditions or cell types (**Supplemental Table S1**). We investigated the use of threshold-free approaches, to allow for recurrent low-affinity binding; and the use of motif-guided signal detection. Overall, we find a surprisingly high concordance of *direct* TF binding across different cell types, tissues and conditions. This is in contrast to previous studies where ChIP-seq peaks are often compared across conditions, yielding poor overlap (Chèneby et al., 2018; Verfaillie et al., 2016). We believe that this is due to the combination of a threshold applied to peak calling, and the mixture of peaks that represent direct TF binding, indirect TF binding, and crosslinking artifacts (i.e., phantom peaks (Baranello et al., 2016; Keren and Segal, 2013). We exploit this discovery to define core sets of preserved direct binding sites to discovery new enhancer features and to train better machine learning models to predict genome-wide TF binding sites. To our knowledge, meta-analysis on ChIP-seq data has not been performed, while meta-analysis of gene expression data is quite common (Brown et al., 2017; Moreau et al., 2003).

After introducing a new TF ChIP-seq meta-analysis approach, based on directly bound regions, and integrated using rank aggregation methods, we showed that the core set of directly bound regions yields high-quality training data for machine learning classifiers. Random Forest models could accurately discriminate bona fide TF binding sites from unbound sites using sequence information only (thus, without using, for example, sequence conservation or chromatin data). Note that this meta-analysis strategy is independent of peak calling, and thereby avoids threshold issues (thresholding otherwise makes cross-data set peak comparisons unfeasible).

The comparison of sequence features of six different TFs revealed several new insights about TF binding modalities, suggesting that these TFs represent different classes (Figure 7). Particularly, “Solitary” TFs, here represented by TP53 and REST, bind to unsophisticated enhancers that harbor a single strong binding site. Interestingly, DHS data analysis supports that these two TFs work alone since their bound regions are associated with the narrowest DHS peaks among the group of studied factors. On the other hand, HIF1A and PPARG represent “Followers”, which are dependent on other factors or features. For these TFs, their motif score is uninformative, and co-factor information is needed and improves the prediction of their binding sites. Finally, SOX2 and GRHL2 represent “Pioneers”, for which GC content and nucleosome occupancy are the strongest features. The finding based on nucleosome occupancy corresponds to previously published studies where SOX2 and GRHL2 are reported as pioneer factors (Jacobs et al., 2018; Soufi et al., 2015).

**Figure 7.**
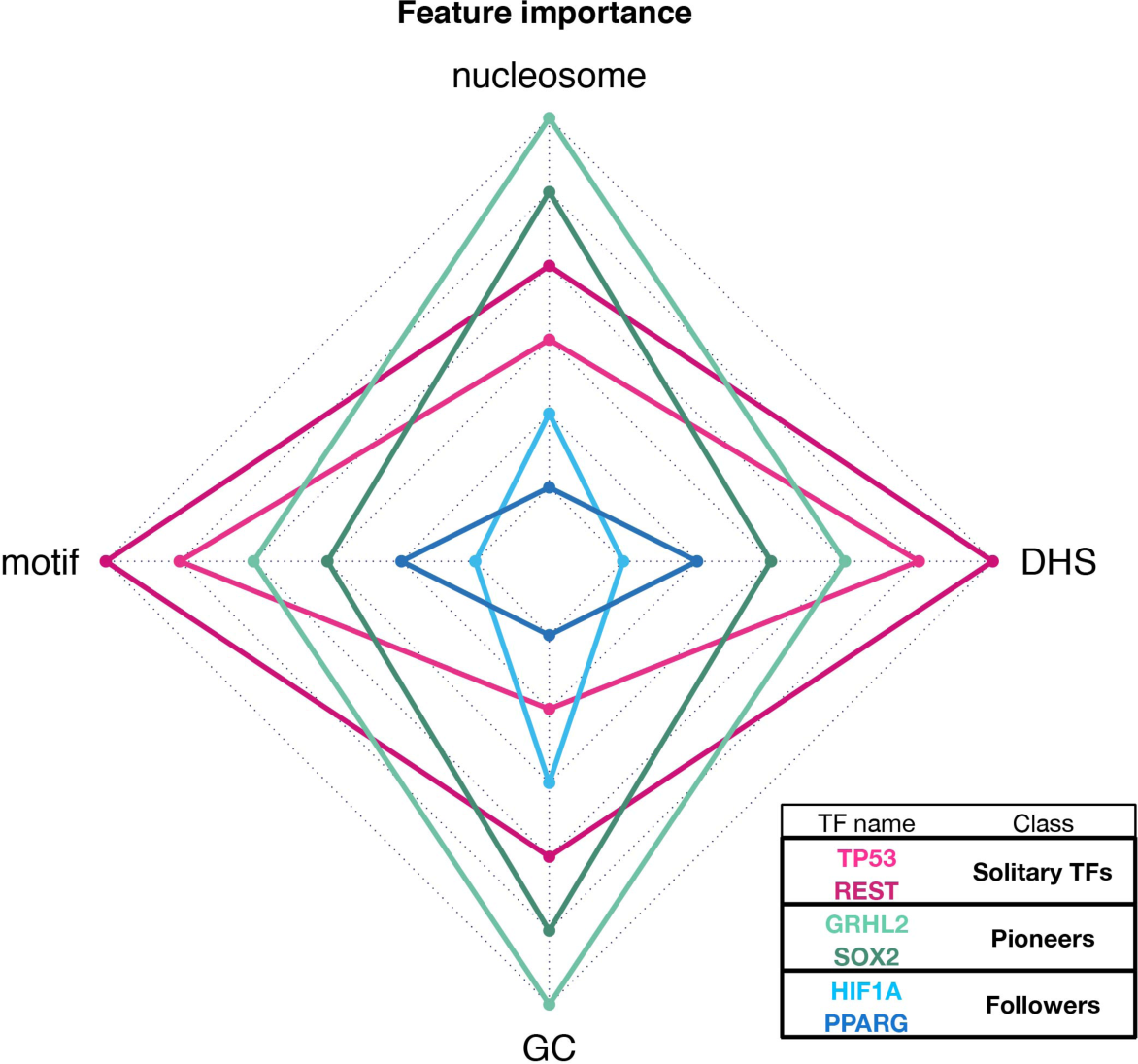
Possible classes of TFs according to selected features, such as preferences to bind nucleosome occupied regions (nucleosome), GC content (GC), differences in motif strength in the human genome computed by cbust (motif), the median of length of DHS peaks overlapping the bound regions (DHS).

In conclusion, to improve the computational prediction of TF binding sites, meta-analysis of ChIP-seq data yields valuable training data and better enhancer models. Depending on the type of TF under study, different types of features are required to train optimal enhancer models.

## Supplementary material

**Supplementary Figure S1.** The table includes logos of PWMs annotated for the specific TFs. These PWMs were used in our study to train Random Forest models using 1 motif (M1 model) and 6 motifs (M6 model).

**Supplementary Figure S2.** Random forest predictions of HIF1A and PPARG binding sites. The plots represent Precision-Recall curves assessing the performance of a Random Forest classifier to discriminate between bound and unbound regions. The performance increased by 10% when PWMs of co-factors were added in the model.

**Supplementary Figure S3.** DNaseI signal on neuronal and neuronal cell lines mapped on the regulatory regions ranked based on REST ChIP-seq meta-analysis.

**Supplementary Table S1.** List of TF ChIP-seq datasets used in the meta-analysis.

**Supplementary Table S2.** i-cisTarget results for TP53 peaks of individual TF ChIP-seq experiments

**Supplementary Table S3.** i-cisTarget results for REST peaks of individual TF ChIP-seq experiments

**Supplementary Table S4.** i-cisTarget results for GRHL2 peaks of individual TF ChIP-seq experiments

**Supplementary Table S5.** i-cisTarget results for SOX2 peaks of individual TF ChIP-seq experiments

**Supplementary Table S6.** i-cisTarget results for HIF1A peaks of individual TF ChIP-seq experiments

**Supplementary Table S7.** i-cisTarget results for PPARG peaks of individual TF ChIP-seq experiments

